# Calibration of in-frame indel variant effect predictors for clinical variant classification

**DOI:** 10.64898/2026.04.15.718599

**Authors:** Haneen Abderrazzaq, Mugdha Singh, Larry Babb, Timothy Bergquist, Steven E. Brenner, Vikas Pejaver, Anne O’Donnell-Luria, Predrag Radivojac, ClinGen Computational Working Group, ClinGen Variant Classification Working Group

## Abstract

Insertions and deletions (indels) represent a substantial source of genetic variation in humans and are associated with a diverse array of functional consequences. Despite their prevalence and clinical importance, indels, particularly short in-frame indels, remain critically understudied compared to single nucleotide variants and are challenging to interpret clinically. While many computational predictors for missense variants have been rigorously evaluated and calibrated for clinical use, the clinical utility of tools for in-frame indels remains uncertain. To address this gap, we have calibrated in-frame indel prediction tools for clinical variant classification. We constructed a high-confidence dataset of in-frame indel variants (≤ 50bp) from clinical and population databases and estimated the prior probability of pathogenicity of a rare in-frame indel observed in a disease-associated gene, and of an insertion and deletion separately. Using a previously developed statistical framework based on local posterior probabilities, we then established score thresholds for eight computational tools, corresponding to distinct evidence levels for pathogenic and benign classification according to ACMG/AMP guidelines. All in-frame indel predictors evaluated here reached multiple evidence levels of pathogenicity and/or benignity, demonstrating measurable clinical value. However, these models consistently exhibited lower performance levels compared to missense predictors, highlighting the need for improved computational approaches for indel classification.

## Introduction

In-frame insertions and deletions (indels) are a common class of genetic variants, exhibiting considerable diversity in their molecular and phenotypic consequences. In contrast to frameshift indels which disrupt the downstream reading frame, typically resulting in premature termination and loss of protein function, in-frame indels preserve the reading frame and result in a protein of altered length while leaving the downstream amino acid sequence intact. The outcomes of in-frame indels are often difficult to predict, as their effects can be more subtle and context-dependent than those of frameshift and nonsense variants.

The abundance of missense variants in variant databases has driven the development of numerous computational models for missense variant effect prediction (Lin et al., 2024). By contrast, the paucity of data on in-frame indels has constrained the development and benchmarking of available computational predictors for this variant class. Cannon et al. (2023) evaluated in-frame indel predictors and reported strong performance across tools on a dataset including ClinVar and gnomAD variants. However, this evaluation did not filter for training data contamination, possibly explaining why tool performance declined substantially on a smaller, novel dataset (Cannon et al., 2023). Additionally, this study applied default or author-recommended classification thresholds rather than thresholds specifically calibrated to clinical evidence standards. To ensure computational tools meet the stringent requirements for clinical use, rigorous calibration using nonoverlapping training and test datasets is essential for establishing appropriate thresholds, but has yet to be done for in-frame indel predictors.

The American College of Medical Genetics and Genomics (ACMG) and the Association for Molecular Pathology (AMP) have published guidelines for clinical variant classification (Richards et al., 2015). These guidelines outline criteria for combining different types of evidence (functional, population, computational, etc.), weighted by strength (supporting, moderate, strong, etc.), to classify a variant as (likely) pathogenic, (likely) benign, or of uncertain significance. Tavtigian et al. (2018, 2020) subsequently developed a Bayesian adaptation of ACMG/AMP guidelines, establishing a more quantitative framework. Given the totality of available evidence, this system requires the posterior probability that a variant is pathogenic to meet thresholds set by the ACMG/AMP guidelines (≥0.99 for pathogenic, ≥0.9 for likely pathogenic, ≤0.1 for likely benign, and ≤0.01 for benign) but specifies that the evidence be quantified in points. In this context, we define calibration as the process of converting evidence such as clinical observations, experimental measurements or prediction scores into additive (with certain exceptions) point values that satisfy the ACMG/AMP posterior probability criteria.

Pejaver et al. (2022) established a framework for aligning computational predictor outputs with the ACMG/AMP guidelines, specifically focusing on missense variant effect predictors. This work calibrated 13 computational tools for missense variants and established the score thresholds for which outputs from each tool correspond to supporting, moderate, or strong evidence (Pejaver et al., 2022). Four of these predictors, along with another three calibrated by Bergquist et al. (2025), achieved strong evidence for pathogenicity, underscoring the growing role of computational predictors in clinical variant classification. Calibrating predictors for in-frame indels can similarly guide laboratories in their use of computational tools, increasing the rigor of variant classification and ultimately improving diagnostic outcomes for patients.

In this work, we apply the Pejaver et al. (2022) framework to in-frame indel predictors to define score thresholds that can be used for clinical variant classification. We estimate the prior probability of pathogenicity for in-frame indels, as well as for insertions and deletions separately, and subsequently define evidence-strength thresholds for these variant types. We evaluate a diverse set of predictors, ranging from traditional models to protein language models. Our results demonstrate that existing in-frame indel tools can be effectively applied for clinical variant classification.

## Materials and Methods

### Datasets

#### ClinVar 2023 calibration set

We extracted in-frame indel variants (insertions, deletions, and complex indel variants) from the December 3, 2023 version of ClinVar (Landrum et al., 2016) based on MANE (Matched Annotation from NCBI and EMBL-EBI) Select transcript annotations (Morales et al., 2022). We maintained both GRCh37 and GRCh38 coordinates for each variant, as prediction tools vary in their accepted genome build input. We further selected variants in disease-associated genes, defined as genes appearing in the Gene Curation Coalition (GenCC) database (DiStefano et al., 2022) with a gene-disease validity classification of definitive, strong, or moderate. We also required the variants to have at least one-star review status with no conflicts, and to be absent from or have a global allele frequency (AF) of ≤ 1% in gnomAD v3.1.2 genomes; genome data were used given their higher indel calling accuracy compared to exome data (Belkadi et al., 2015). Variants were additionally restricted to ≤ 50 bp in length to exclude structural variants. Finally, only variants with one of the following classifications were retained: pathogenic, pathogenic/likely pathogenic, likely pathogenic, benign, benign/likely benign, or likely benign, with no conflicting assertions allowed. The final dataset comprised 3,625 indels, including 1,979 pathogenic/likely pathogenic and 1,646 benign/likely benign variants, with length distributions shown in Figure 1A-B. These variants were distributed across 1,113 disease-associated genes (Supplementary Figure 1). For analyses comparing insertions and deletions, complex indel variants were classified based on their net effect on sequence length change. The ClinVar 2023 dataset is available in Supplementary File 1.

**Figure 1.**
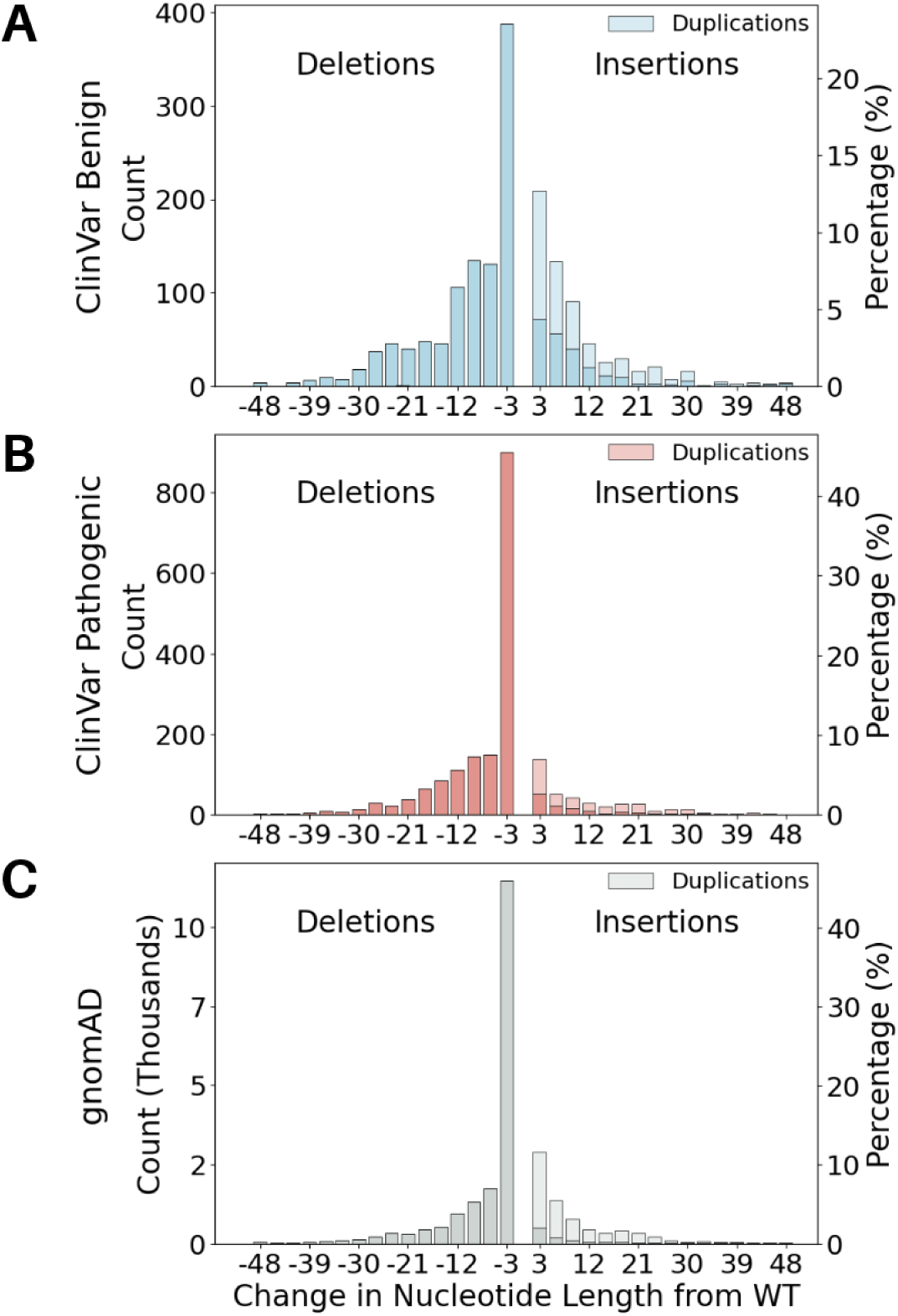
Length distribution of the indels in the ClinVar 2023 (A-B) and gnomAD (C) datasets, separated into variants labeled as benign (A) and pathogenic (B). Deletions are considered to have a negative change in nucleotide length from the wild-type sequence, and insertions have a positive change. Duplications, a subset of insertions, are shown in a lighter color on each graph.

#### ClinVar 2025 test set

We created a separate dataset of ClinVar variants to evaluate our thresholds using the December 1, 2025 version of ClinVar, applying the same filtering steps described above for the ClinVar 2023 dataset. Any variants present in the 2023 set were excluded. The resulting test dataset contained 1,131 indels, including 793 pathogenic/likely pathogenic and 338 benign/likely benign variants, distributed across 559 disease-associated genes (Supplementary Figure 2). The ClinVar 2025 dataset is available in Supplementary File 1.

#### gnomAD set

We also constructed a dataset of rare, naturally occurring indel variants from gnomAD (Karczewski et al., 2020) to serve as a reference set of human in-frame indel variation. We used gnomAD v2.1.1 exomes and genomes, which is mapped to the GRCh37 reference genome build required for input to some of our tools. We selected variants in genes appearing in GenCC (definitive, strong, or moderate classification), with a global AF ≤ 1%, classified as in-frame based on the MANE Select transcript (v1.5), ≤ 50 bp in length, and passing all variant quality filters. This yielded a final dataset of 26,014 variants, the length distribution of which is shown in Figure 1C. The gnomAD dataset is available in Supplementary File 1.

#### Rare Genomes Project set

We also constructed a rare disease dataset from the Rare Genomes Project (RGP) (Serrano et al., 2023) to determine the extent of in-frame indel variation and effects of calibration when considering an individual’s genome. The 300 RGP probands included here were previously used to assess missense calibration (Stenton et al., 2024). Variants were called with GATK version 4.1.8.0 on GRCh38 (McKenna et al., 2010). We selected variants with genotype quality ≥ 40, depth ≥ 10, allele balance ≥ 0.2, and indel size ≤ 50 bp. MANE Select transcripts with “inframe_deletion” and “inframe_insertion” consequences were retained. To ensure relevance for rare disease analyses, we further selected variants that were either absent from or have a global AF ≤ 1% in gnomAD v3.1.2 genomes, and filtered for disease-associated genes with definitive, strong, or moderate evidence in GenCC. The subjects from the RGP cohort were distinct from those used to construct gnomAD and had signed informed consent to use the data for research purposes (Mass General Brigham IRB protocol 2016P001422).

### Selection of Computational Tools

The tools selected for analysis were CADD v1.7 (Schubach et al., 2024), ESM1b (Rives et al., 2021), FATHMM-indel (Ferlaino et al., 2017), INDELpred (Wei et al., 2024), MutPred-Indel (Pagel et al., 2019), ProGen2 (Nijkamp et al., 2023), PROVEAN (Choi et al., 2012) and VEST-Indel (Douville et al., 2016). We included tools based on clinical community interest, user accessibility, and availability of training data (directly or after communicating with the developers), for exclusion from our calibration dataset where necessary. As in previous work, tools that use allele frequency as a feature were omitted (Pejaver et al., 2022). We also sought to evaluate a diverse set of methods, including evolutionary conservation-based methods such as PROVEAN, traditional machine learning models such as VEST-Indel and MutPred-Indel, and protein language models such as ESM1b and ProGen2. Protein language models refer to neural networks that treat amino acid sequences as sentences and identify patterns in sequence data by adapting techniques developed for natural language processing. In contrast to traditional methods, these models are typically trained without class labels to learn representations (vector embeddings) of protein sequences, though they implicitly capture evolutionary information.

### Training data filtering

Training datasets were obtained for all tools built in a supervised manner (MutPred-Indel, VEST-Indel, FATHMM-indel, and INDELpred) and any training variants overlapping with the ClinVar 2023, ClinVar 2025, or gnomAD sets were filtered out for each tool separately. This was done to avoid biased performance assessment, as predictors inherently perform better on data encountered in training. The number of variants remaining for each tool after this filtering step can be found in Supplementary Table 1. After removal of training data, between 1,783 (INDELpred) and 3,625 (CADD, ESM1b, ProGen2, and PROVEAN) ClinVar 2023 variants for use in calibration.

### Score Generation

Some tools only accept GRCh37 coordinates, necessitating the use of both genome builds. For ClinVar data, GRCh37 variant coordinates were used for FATHMM-indel, INDELpred, and PROVEAN, while GRCh38 coordinates were used for VEST-Indel and CADD. GRCh37 data were used for all these tools for gnomAD and RGP data, with RGP coordinates lifted over from GRCh38 to GRCh37 using PyLiftover with UCSC chain files. For MutPred-Indel, ESM1b, and ProGen2, protein sequence input was generated from GRCh38 data for ClinVar variants and GRCh37 data for gnomAD and RGP variants using the June 2020 release of ANNOVAR (Wang et al., 2010), with annotations restricted to MANE Select transcripts.

While precomputed tables of single nucleotide variants are available for many tools (Liu et al., 2020), there are a limited number of indels scored in these files. Thus, relevant prediction scores were generated for this project. For VEST-Indel, FATHMM-indel, CADD, and PROVEAN, scores were generated using each tool’s web server; see Supplementary Materials for URLs to each portal. We used the condensed results for PROVEAN. For the remainder of the tools, the scores were generated using available source code, even when a web server was also available. Unlike the other models, ESM1b and ProGen2 do not directly provide deleteriousness predictions; we instead used the log-likelihood ratio of the variant protein sequence compared to the wild-type sequence as a proxy for a prediction score. A variant sequence with a lower likelihood score than the corresponding wild-type sequence is, in principle, less likely to represent a functional protein, and thus the variant is more likely to be deleterious.

For ESM1b, we used the workflow from Brandes et al. (2023), which directly outputs log-likelihood ratios for wild-type versus variant protein sequences. For ProGen2 (base model), we obtained log-likelihood scores for the wild-type and variant sequences independently and calculated the log-likelihood ratios as log-likelihood of the variant sequence minus that of the wild-type sequence. ProGen2 limits input protein sequences to a maximum length of 2,048 amino acids, including the start and end tokens. As such, sequences in our dataset longer than 2,046 amino acids were cropped prior to input into the tool. Specifically, a window of length 2,046 amino acids was centered around the midpoint of the start and end coordinates of the variant position and applied to both the wild-type and variant protein sequences. If centering this window would extend beyond the protein N- or C-terminus, the window was shifted to stay within protein boundaries.

The pairwise correlation between the outputs of all evaluated tools on the gnomAD set is presented in Supplementary Figure 3.

### Calibration methodology

The prior probability of pathogenicity for missense variants was previously established as 4.4% (Pejaver et al., 2022). We applied the same framework to in-frame indels, using the gnomAD set described above as the reference set and the pathogenic variants from the ClinVar 2023 set as the positive set. Overlapping variants were not removed from gnomAD for the prior estimation process. We used features generated using MutPred-Indel and applied the DistCurve algorithm (Zeiberg et al., 2020) to estimate the prior for all in-frame indels, as well as for insertions and deletions separately. For the purposes of calibration, we removed 1,078 variants from the gnomAD dataset that were present in the ClinVar 2023 calibration set (863 benign, 215 pathogenic) to yield 24,936 indels. We then performed calibration for insertions and deletions independently, applying the local posterior probability framework according to Pejaver et al. (2022). Thresholds lacking sufficient data support at the extremes of the score distribution were subsequently excluded.

## Results

### In-frame variation in an individual’s genome

In assessing the 300 probands from the Rare Genomes Project (RGP), we observed that each individual harbored, on average, 8,900 missense variants (range: 8,383-10,616), 310 in-frame indels (range: 260-372), 112 frameshift variants (range: 89-150), and 58 nonsense variants (range: 43-79) (Figure 2A). When filtered for rare variants (absent from gnomAD or AF ≤ 1% in gnomAD v3.1.2 genomes), each proband harbored a mean of 470 missense (range: 323-1,229), 15 in-frame indels (range: 4-39), 9 frameshift variants (range: 2-25), and 7 nonsense (range: 2-20) rare variants (Figure 2B). For in-frame indels specifically, we further assessed rare variants in disease-associated genes (definitive, strong, or moderate evidence in GenCC). Nearly all probands (293 out of 300, 97%) harbored rare in-frame indels in disease-associated genes, with a mean of approximately 4 (range: 1-10) per individual (Figure 2C). When stratified by variant type, these rare indels comprised a mean of approximately 1 in-frame insertion and 2 in-frame deletions per proband (Figure 2D), demonstrating that, despite being less frequent than missense variants, they are sufficiently prevalent to warrant well-calibrated computational prediction tools for clinical variant classification.

**Figure 2.**
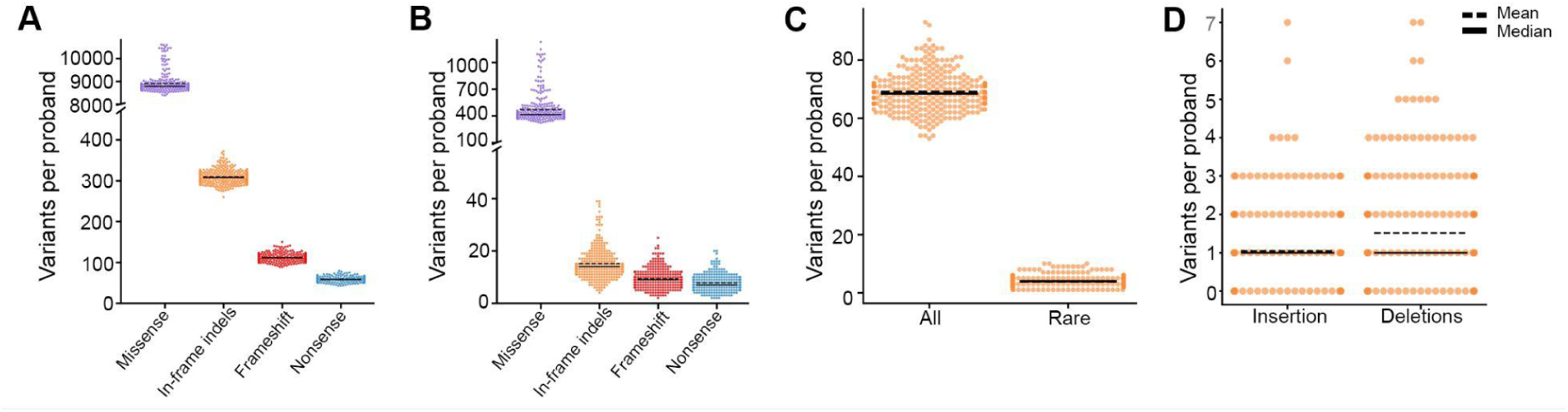
(A) Total number of missense (purple), in-frame indel (orange), frameshift (red) and nonsense variants (blue) per proband. (B) Rare (AF ≤ 1%) missense, in-frame indel, frameshift and nonsense variants per proband. (C) Total and rare in-frame indels in disease-associated genes per proband and (D) number of rare in-frame insertions and deletions in disease associated genes.

### Estimating prior probability of pathogenicity

Using gnomAD as the reference set of human variation, the prior probability of pathogenicity for short (≤ 50 bp) in-frame indels was estimated to be 4.0%, slightly lower than the 4.4% previously estimated for missense variants (Pejaver et al., 2022). When separating insertions and deletions, we found that insertions (0.8%) had a substantially lower prior probability than deletions (4.6%), with the latter being approximately equivalent to that of missense variants (Pejaver et al., 2022). This suggests that in-frame insertions are less likely to be pathogenic than in-frame deletions, which is also supported by the data in Figure 1 where 38.9% of insertions (392 out of 1,009) in our ClinVar 2023 calibration set were labeled as pathogenic compared to 60.7% of deletions (1,587 out of 2,616). This prior estimation relies on an assumption that the pathogenic variants used for estimation were representative of all pathogenic variants for both insertions and deletions (Zeiberg et al., 2020).

### Calibrated tools often reach -1 to +2 strength of evidence

The estimated posterior probability curves for all eight tools are shown in Figure 3 for insertions and deletions separately. From these curves, we determined score thresholds for all tools and levels of evidence strength for insertions and deletions independently (Tables 1 and 2). We report thresholds for up to four points of evidence in both directions for each tool, although PROVEAN and INDELpred reached higher evidence levels for benignity. No tools reached strong evidence of pathogenicity; the highest pathogenic evidence level achieved was +3, reached by MutPred-Indel, VEST-Indel, ESM1b, ProGen2, and PROVEAN for deletions, and by VEST-Indel for insertions. FATHMM-indel, INDELpred, and PROVEAN reached strong levels for benignity (-4) for deletions and INDELpred did the same for insertions. All predictors reach the supporting level of benignity (-1) for both insertions and deletions, with the exception of ProGen2, which reaches it for deletions only, and ESM1b which does not reach it for either variant type. The variant counts per bin for each met evidence strength level are reported in Supplementary Figure 4.

**Figure 3.**
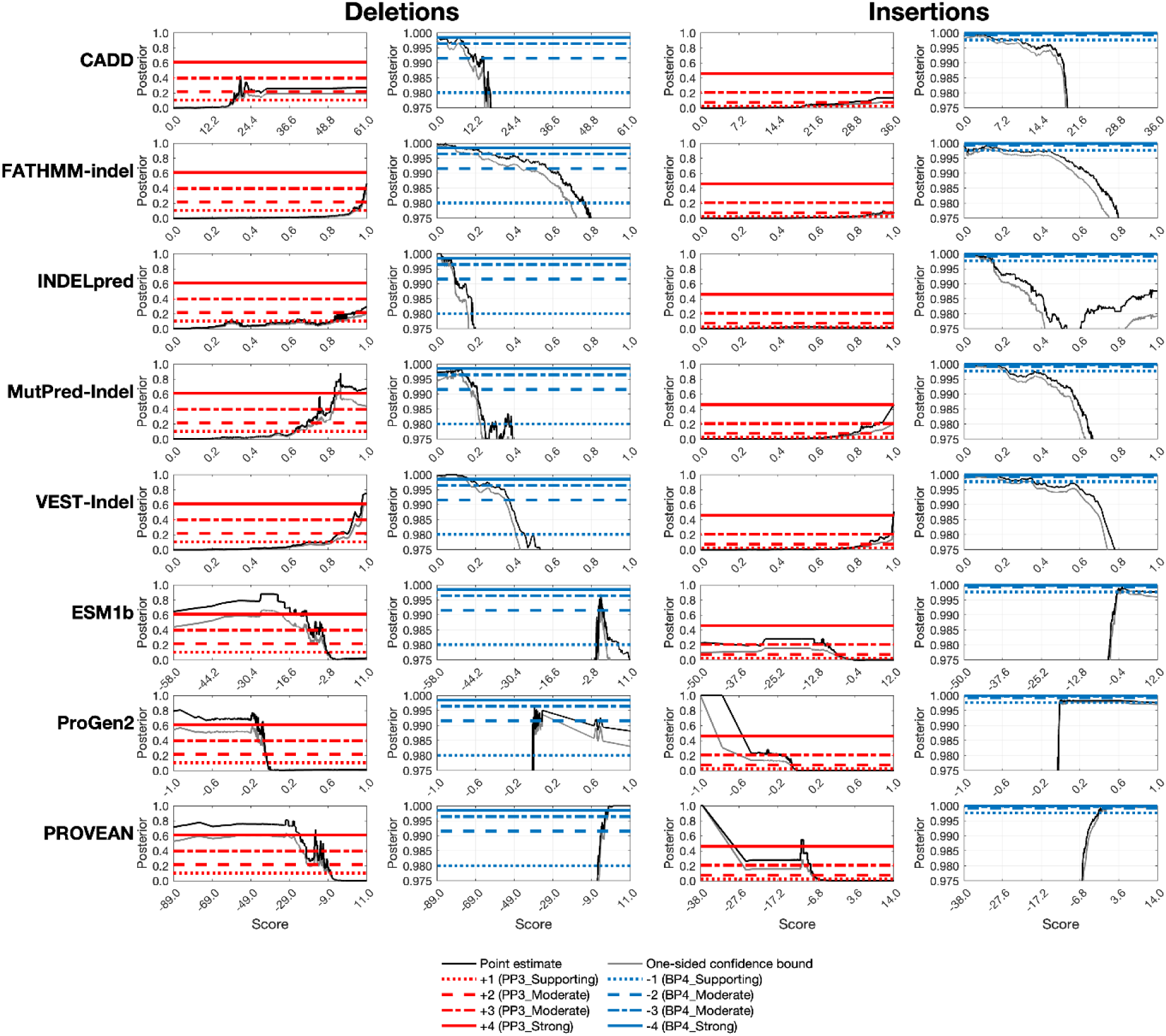
Final local posterior probability curves for each tool for insertions and deletions, using the estimated prior probabilities of 0.8% and 4.6% respectively. The horizontal lines represent four different evidence point values: +1, +2, +3, and +4 for pathogenic curves and -1, -2, -3, and -4 for benign curves. The black curve is the local posterior probability estimate, and the grey curve represents the one-sided 95% confidence interval (on the more stringent side) after 10,000 bootstrap iterations. The first five tools shown use lower scores to mean that a variant is more likely benign and higher scores to mean that a variant is more likely pathogenic; the last three tools do the opposite. Within these groups, tools are sorted alphabetically.

Predictors generally achieved higher evidence levels for deletions than for insertions, a disparity that likely reflects the lower prior probability of pathogenicity for insertions (an effect shown by The Critical Assessment of Genome Interpretation Consortium, 2024; Figure 3 therein). The disparity likely also reflects the limited representation of pathogenic insertions in predictor training datasets and the difficulty of evaluating a new sequence introduced into an existing protein without any functional domain evidence. Differences in both the maximum evidence levels achieved and the thresholds required to reach equivalent evidence levels underscored the necessity of calibrating insertions and deletions separately.

In several cases, the thresholds (Tables 1 and 2) for reaching even the lowest level of pathogenic evidence (+1) diverged significantly from the default or author-recommended thresholds. For FATHMM-indel, our calibrated +1 thresholds were 0.961 for deletions and 0.845 for insertions, which are considerably higher than the tool’s default classification threshold of 0.5. Similarly, -2.5 was the recommended classification threshold for PROVEAN, with scores below -2.5 being classified as pathogenic. However, the established thresholds for the +1 evidence level are -9.10 for deletions and -6.69 for insertions. These discrepancies demonstrate the importance of using calibrated thresholds for variant classification; applying default thresholds for these tools would likely result in substantial overweighting of evidence.

**Table 1.**
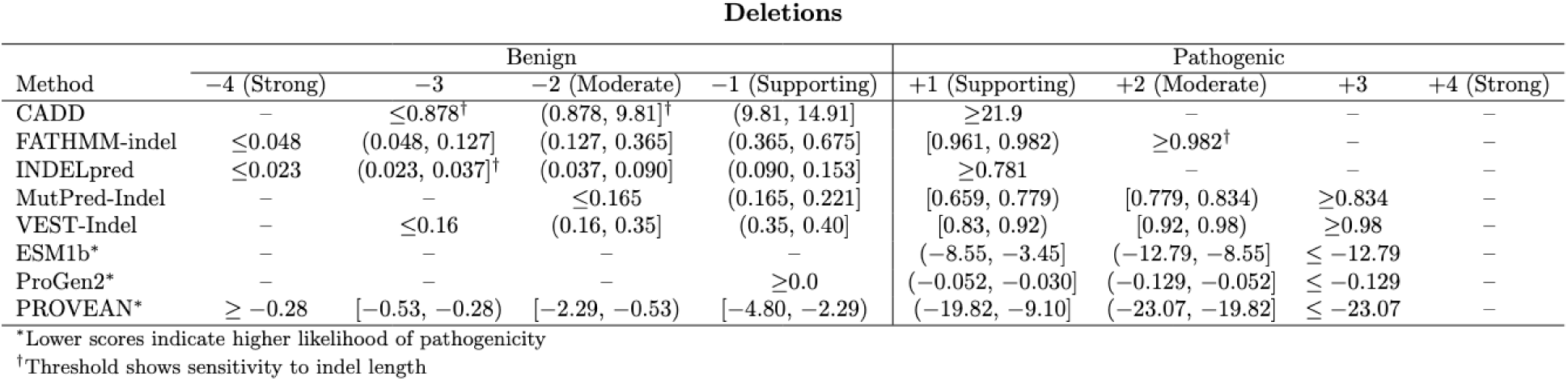
Final score thresholds for each tool for deletions. For CADD, FATHMM-indel, INDELpred, MutPred-Indel, and VEST-Indel, higher scores indicate a higher likelihood of pathogenicity. For ESM1b, PROVEAN, and ProGen2, lower scores indicate a higher likelihood of pathogenicity. Several tools showed some sensitivity to indel length (Supplementary Figure 6) but current datasets are too small for length-based calibration (Supplementary Table 3).

**Table 2.** Final score thresholds for each tool for insertions. As with deletions, CADD, FATHMM-indel, INDELpred, MutPred-Indel, and VEST-Indel consider higher scores to indicate higher likelihood of pathogenicity while lower scores are indicative for ESM1b, PROVEAN, and ProGen2. Several tools showed some sensitivity to indel length (Supplementary Figure 6) but current datasets are too small for length-based calibration (Supplementary Table 3).

| Insertions |  |  |  |  |  |  |  |  |
| --- | --- | --- | --- | --- | --- | --- | --- | --- |
| Method | Benign |  |  |  | Pathogenic |  |  |  |
|  | -4 (Strong) | -3 | -2 (Moderate) | -1 (Supporting) | +1 (Supporting) | +2 (Moderate) | +3 | +4 (Strong) |
| CADD | - | - | - | ≤4.72 | [20.3, 34.0)<sup>†</sup> | ≥34.0 | - | - |
| FATHMM-indel | - | - | - | ≤0.008 | ≥0.845 | - | - | - |
| INDELpred | ≤0.008 | (0.008, 0.010] | (0.010, 0.054]<sup>†</sup> | (0.054, 0.127] | - | - | - | - |
| MutPred-Indel | - | - | - | ≤0.212 | [0.691, 0.838) | ≥0.838 | - | - |
| VEST-Indel | - | - | - | ≤0.21 | [0.80, 0.91) | [0.91, 1.0)<sup>†</sup> | 1.0 | - |
| ESM1b* | - | - | - | - | (-7.52, -4.25] | ≤-7.52 | - | - |
| ProGen2* | - | - | - | - | (-0.076, -0.037] | ≤-0.076 | - | - |
| PROVEAN* | - | - | ≥0.64 | [-1.19, 0.64) | (-8.17, -6.69] | ≤-8.17 | - | - |

\*Lower scores indicate higher likelihood of pathogenicity
$^\dagger$ Threshold shows sensitivity to indel length

### Evaluation of proposed thresholds

For each predictor we calculated the percentage of gnomAD variants that would be assigned each level of evidence given the newly estimated thresholds (Table 3). Predictors varied considerably in their ability to assign evidence. PROVEAN had the highest percentages, assigning 76.9% of deletions and 55.6% of insertions at least supporting evidence for either pathogenicity or benignity, while INDELpred assigned evidence to only 12.3% of deletions and 13.0% of insertions. Note, however, that INDELpred had the highest fraction of variants removed for calibration due to overlap with its training set, leading to higher statistical uncertainty when establishing thresholds and correspondingly stricter evidence thresholds. Across all predictors that reached +3 evidence, only a small proportion of gnomAD variants fell into these bins, which gives confidence that the newly established thresholds would not lead to overprediction of pathogenic variants.

**Table 3.** Percentage of variants in the gnomAD set falling into each bin given calibrated thresholds (Tables 1 and 2) for deletions (left) and insertions (right).

| Deletions |  |  |  |  |  |  |  |  |  |
| --- | --- | --- | --- | --- | --- | --- | --- | --- | --- |
|  | -4 | -3 | -2 | -1 | Indet. | +1 | +2 | +3 | +4 |
| CADD |  | 0.3 | 6.7 | 12.0 | 74.0 | 7.0 |  |  |  |
| FATHMM-Indel | 5.2 | 4.2 | 7.0 | 8.8 | 35.3 | 17.2 | 22.3 |  |  |
| INDELpred | 3.7 | 0.7 | 4.5 | 2.7 | 87.7 | 0.5 |  |  |  |
| MutPred-Indel |  |  | 15.0 | 9.6 | 58.3 | 10.4 | 3.3 | 3.4 |  |
| VEST-Indel |  | 13.7 | 16.6 | 3.7 | 36.6 | 11.3 | 10.3 | 7.7 |  |
| ESM1b |  |  |  |  | 75.2 | 15.4 | 5.6 | 3.7 |  |
| ProGen2 |  |  |  | 25.1 | 45.0 | 12.0 | 13.8 | 4.1 |  |
| PROVEAN | 8.5 | 1.8 | 15.3 | 19.6 | 23.1 | 27.2 | 1.3 | 3.1 |  |

| Insertions |  |  |  |  |  |  |  |  |  |
| --- | --- | --- | --- | --- | --- | --- | --- | --- | --- |
|  | -4 | -3 | -2 | -1 | Indet. | +1 | +2 | +3 | +4 |
| CADD |  |  |  | 9.1 | 77.2 | 13.6 | 0.0 |  |  |
| FATHMM-Indel |  |  |  | 4.2 | 62.0 | 33.7 |  |  |  |
| INDELpred | 1.2 | 0.4 | 7.4 | 4.0 | 87.0 |  |  |  |  |
| MutPred-Indel |  |  |  | 25.1 | 62.3 | 10.2 | 2.4 |  |  |
| VEST-Indel |  |  |  | 25.9 | 51.6 | 9.7 | 11.2 | 1.7 |  |
| ESM1b |  |  |  |  | 83.1 | 8.2 | 8.8 |  |  |
| ProGen2 |  |  |  |  | 81.5 | 11.4 | 7.1 |  |  |
| PROVEAN |  |  | 5.5 | 25.9 | 44.4 | 6.8 | 17.5 |  |  |

We also calculated the positive likelihood ratios (LRs) on the ClinVar 2025 set, which served as an independent test set, to determine how well the new thresholds applied to unseen variants (Table 4). Based on the framework proposed by Tavtigian et al. (2018), which defined the minimum positive likelihood ratio expected at each evidence level given the prior probability of pathogenicity, we expected likelihood ratios of 2.39 (supporting, +1), 5.69 (moderate, +2), 13.59 (moderate, +3) and 32.42 (strong, +4) for deletions and 3.25 (supporting, +1), 10.55 (moderate, +2), 34.28 (moderate, +3) and 111.34 (strong, +4) for insertions. For each tool, all achieved evidence levels met the corresponding likelihood ratio requirements, except for CADD and INDELpred at the -3 deletion threshold, though the observed ratios remained high. This analysis was limited by the size of the dataset due to the relatively small number of in-frame indels, especially insertions, that have accumulated in ClinVar over the past two years. The variant counts per bin for this dataset are reported in Supplementary Figure 5.

We also sought to assess threshold performance across indels of different lengths. We calculated LRs on our ClinVar 2025 dataset separately for single amino acid indels versus longer indels (Supplementary Figure 6), following the same approach as Table 4. Most tools met the expected LRs for both single amino acid and multiple amino acid insertions and deletions. The few exceptions likely reflect limited data, as small numbers of misclassifications can substantially affect LR estimates. In general, our calibration was robust to indel length, although caution is advised when using CADD and INDELpred, as both failed to meet expected LRs across multiple length-stratified bins. At the time of this analysis, there was not enough data in each of these categories to have confidently performed prior probability estimation and calibrations for indels separated by length (Supplementary Table 2). Length-specific calibration may be explored in the future once sufficient data accumulate.

**Table 4.** Likelihood ratios calculated on ClinVar 2025 set for deletions (left) and insertions (right). For benign evidence (blue), inverse likelihood ratios (FPR/TPR) are shown. For pathogenic evidence (red), standard likelihood ratios (TPR/FPR) are shown. On both sides, higher values indicate stronger classification performance on the test data. An infinite value indicates that there were no incorrect predictions in that bin. The expected likelihood ratios for deletions are 2.39 (supporting, +1), 5.69 (moderate, +2), 13.59 (moderate, +3) and 32.42 (strong, +4). The expected likelihood ratios for insertions are 3.25 (supporting, +1), 10.55 (moderate, +2), 34.28 (moderate, +3) and 111.34 (strong, +4).

| Deletions |  |  |  |  |  |  |  |  |  |
| --- | --- | --- | --- | --- | --- | --- | --- | --- | --- |
| CADD |  | 12.2 | 5.7 | 4.9 |  | 5.4 |  |  |  |
| FATHMM-indel | 57.5 | 21.6 | 18.1 | 6.1 |  | 3.2 | 8.7 |  |  |
| INDELpred | 59.6 | 6.0 | 17.9 | 11.9 |  | 8.1 |  |  |  |
| MutPred-Indel |  |  | 33.2 | 10.0 |  | 5.3 | 25.7 | ∞ |  |
| VEST-Indel |  | 30.0 | 37.5 | 9.8 |  | 6.1 | 8.9 | 32.7 |  |
| ESM1b |  |  |  |  |  | 21.9 | ∞ | ∞ |  |
| ProGen2 |  |  |  | 3.9 |  | 8.6 | 11.2 | ∞ |  |
| PROVEAN | ∞ | 39.2 | 23.7 | 5.9 |  | 6.9 | 14.3 | 57.1 |  |
|  | -4 | -3 | -2 | -1 |  | +1 | +2 | +3 | +4 |

| Insertions |  |  |  |  |  |  |  |  |  |
| --- | --- | --- | --- | --- | --- | --- | --- | --- | --- |
| CADD |  |  |  | 5.8 |  | 3.4 | ∞ |  |  |
| FATHMM-indel |  |  |  | ∞ |  | 5.1 |  |  |  |
| INDELpred | ∞ | ∞ | 13.8 | ∞ |  |  |  |  |  |
| MutPred-Indel |  |  |  | 18.0 |  | 5.7 | ∞ |  |  |
| VEST-Indel |  |  |  | 39.1 |  | 5.9 | 21.4 | ∞ |  |
| ESM1b |  |  |  |  |  | 7.7 | 23.1 |  |  |
| ProGen2 |  |  |  |  |  | 11.1 | 51.8 |  |  |
| PROVEAN |  |  | ∞ | 13.1 |  | ∞ | 26.8 |  |  |
|  | -4 | -3 | -2 | -1 |  | +1 | +2 | +3 | +4 |

### Evidence yield on the RGP cohort

We evaluated the distribution of rare in-frame indels per RGP proband in disease-associated genes across ACMG/AMP evidence categories with our eight newly calibrated computational prediction tools (Table 5A-B). Unlike gnomAD, RGP provides per-proband variant data, allowing us to estimate the number of variants that our new thresholds will contribute to classification for an individual in a clinical setting, where we intend these calibrated thresholds to be used.

CADD, FATHMM-indel, and INDELpred scored nearly all rare in-frame indels, and PROVEAN and VEST-Indel scored more than 95% of the input variants. MutPred-Indel, ESM1b, and ProGen2 showed a 6% reduction in scored variants, as ANNOVAR was unable to generate the protein sequences required as model input for these variants (Supplementary Table 3).

Across the RGP cohort, approximately four in-frame rare indels per individual in disease-associated genes were observed (Figure 2), which were subsequently resolved into ACMG/AMP categories based on calibrated scores per tool. For each tool, 0-1 indels per proband received benign evidence and 1-3 indels per proband fell into the indeterminate range. A small number of variants per proband received pathogenic evidence from at least one tool; e.g., approximately 1 in 10 probands harbored at least one indel in the +1 pathogenic evidence category (Table 5A-B). Furthermore, approximately 1 in 25 probands were assigned +2 or +3 points of pathogenic evidence by at least one tool; notably, no insertions reached this category for CADD, FATHMM-indel, INDELpred, or PROVEAN and no deletions reached this category for INDELpred (Table 5A-B).

**Table 5.** Number of rare in-frame deletions (A) and in-frame insertions (B) per proband in disease-associated genes across 300 RGP probands for each evidence strength.

| Tool | Benign |  |  |  | Indeterminate | Pathogenic |  |  |
| --- | --- | --- | --- | --- | --- | --- | --- | --- |
|  | -4 | -3 | -2 | -1 |  | +1 | +2 | +3 |
| CADD | – | 0.02 (0-1) | 0.55 (0-4) | 0.50 (0-3) | 1.21 (0-6) | 0.05 (0-1) | – | – |
| FATHMM-indel | 0.61 (0-4) | 0.26 (0-4) | 0.45 (0-4) | 0.24 (0-2) | 0.39 (0-3) | 0.14 (0-2) | 0.20 (0-2) | – |
| INDELpred | 1.47 (0-6) | 0.07 (0-1) | 0.14 (0-2) | 0.05 (0-2) | 0.57 (0-3) | 0.01 (0-1) | – | – |
| MutPred-Indel | – | – | 0.51 (0-3) | 0.16 (0-2) | 1.33 (0-6) | 0.11 (0-2) | 0.03 (0-1) | 0.04 (0-1) |
| VEST-Indel | – | 0.76 (0-5) | 0.77 (0-4) | 0.06 (0-2) | 0.50 (0-5) | 0.11 (0-1) | 0.05 (0-2) | 0.04 (0-1) |
| ESM1b | – | – | – | – | 1.99 (0-7) | 0.14 (0-2) | 0.03 (0-1) | 0.01 (0-1) |
| ProGen2 | – | – | – | 0.70 (0-3) | 1.28 (0-6) | 0.10 (0-2) | 0.08 (0-1) | 0.03 (0-1) |
| PROVEAN | 0.73 (0-4) | 0.06 (0-1) | 0.41 (0-4) | 0.37 (0-4) | 0.33 (0-4) | 0.30 (0-3) | 0.0 (0-1) | 0.04 (0-1) |

| Tool | Benign |  |  |  | Indeterminate | Pathogenic |  |  |
| --- | --- | --- | --- | --- | --- | --- | --- | --- |
|  | -4 | -3 | -2 | -1 |  | +1 | +2 | +3 |
| CADD | – | – | – | 0.29 (0-2) | 1.23 (0-7) | 0.06 (0-1) | – | – |
| FATHMM-indel | – | – | – | 0.11 (0-1) | 1.34 (0-7) | 0.10 (0-2) | – | – |
| INDELpred | 0.61 (0-4) | 0.04 (0-1) | 0.52 (0-4) | 0.07 (0-2) | 0.32 (0-3) | – | – | – |
| MutPred-Indel | – | – | – | 0.51 (0-3) | 0.83 (0-7) | 0.06 (0-2) | 0.01 (0-1) | – |
| VEST-Indel | – | – | – | 0.72 (0-4) | 0.76 (0-5) | 0.05 (0-1) | 0.02 (0-2) | – |
| ESM1b | – | – | – | – | 1.36 (0-7) | 0.03 (0-2) | 0.02 (0-1) | – |
| ProGen2 | – | – | – | – | 1.37 (0-7) | 0.03 (0-2) | 0.01 (0-1) | – |
| PROVEAN | – | – | 0.13 (0-1) | 0.70 (0-4) | 0.58 (0-4) | 0.04 (0-1) | 0.08 (0-2) | – |

## Discussion

We found that in-frame indel variant effect predictors can be successfully calibrated and used to generate evidence for clinical variant classification. All eight predictors assessed met the requirements to provide at least supporting level of evidence for pathogenicity, benignity, or both, across insertions and deletions, and in most cases reached moderate evidence. We applied our calibrated thresholds to several independent datasets (Tables 3, 4, and 5), confirming that these thresholds generalize well without leading to overprediction.

Despite being an older and simpler method, PROVEAN matches the performance of newer tools in terms of the evidence levels it achieved on both the pathogenic and benign sides for insertions and deletions. This finding underscores the value of conservation data in pathogenicity prediction, as previously demonstrated for missense variation (Capriotti & Fariselli, 2022). However, we were unable to find a working software, and researchers and diagnostic labs may therefore be limited to the web portal for its use. The protein language models, ESM1b and ProGen2, had some of the most imbalanced calibration results, performing well on the pathogenic side but poorly on the benign side; true pathogenic variants were often the only ones to receive negative log-likelihood ratios, while both benign and pathogenic variants tended to score near zero, making it difficult to assign benign evidence. Among the other machine learning methods, we observed similar results with more balanced performance for MutPred-Indel and VEST-Indel, as well as FATHMM-indel and INDELpred.

In contrast to these in-frame indel predictor results, several missense predictors have previously been shown to reach strong evidence levels upon calibration (Pejaver et al. 2022; Bergquist et al., 2025): BayesDel (Feng et al., 2017), MutPred2 (Pejaver et al., 2020), REVEL (Ioannidis et al., 2016), VEST4 (Carter et al., 2013), ESM1b (Rives et al., 2021), AlphaMissense (Cheng et al., 2023) and VARITY (Wu et al., 2021). The indel-equivalent models MutPred-Indel and VEST-Indel reached only +3 evidence for deletions, and +2 and +3 for insertions, respectively. CADD, which was calibrated here and by Pejaver et al. (2022), reached the +2 level for both missense variants and insertions, but only +1 evidence for deletions. Collectively, these results suggest that indel predictors still fall behind missense predictors in calibrated evidence strength.

### Recommendations for updates to PP3/BP4 criteria

Tools were previously recommended for clinical use for missense variant classification if they could reach at least strong evidence on the pathogenic side and moderate evidence on the benign side (Pejaver et al., 2022). None of the in-frame indel predictors evaluated here meet this standard. However, three in-frame indel predictors (MutPred-Indel, VEST-Indel, and PROVEAN) reached at least moderate evidence (+2 or +3) for pathogenicity and at least supporting evidence for benignity for both insertions and deletions independently. Other tools showed imbalanced performance, usually reaching +3 or -3 evidence but not both. We recommend selecting a tool that reaches at least moderate pathogenic evidence and supporting benign evidence.

The thresholds established here provide actionable guidance for applying computational tools in clinical variant classification. When selecting a tool, we further recommend selecting a single predictor before variant scoring, consistent with Pejaver et al. (2022) recommendations, since tools were calibrated individually and their joint application has not been evaluated. Laboratories may choose separate tools for insertions and deletions, as some predictors demonstrate superior performance for one variant type. For combining PP3/BP4 evidence with other ACMG/AMP evidence codes, we propose following the recommendations established in Pejaver et al. (2022). We also note that the thresholds established here apply to the specified versions of these tools and may not be valid for future versions of the tools.

While our calibration was performed on genome-wide data, given gene-specific or domain-specific evidence, it is appropriate for Variant Curation Expert Panels (VCEPs) to use alternative thresholds tailored to their genes or domains of interest if sufficient data are available for calibration. Finally, an additional consideration for tool selection is practical accessibility; the ability to score variants on a web server without the requirement of software installation may be preferred in some clinical settings. Web servers are available for CADD, VEST-Indel, FATHMM-indel, PROVEAN, and MutPred-Indel, though query size limits vary by tool.

### Limitations and future work

One limitation of this work is that each tool was evaluated on a different dataset, as training variants were filtered out individually for each tool. This was done to maximize the number of variants available for calibration and obtain more accurate thresholds. INDELpred, which was trained on ClinVar, had approximately half of our ClinVar 2023 variants present in its training set, resulting in a dataset of 1,688 variants to be used for calibration, compared to over 3,000 for most other tools (Supplementary Table 1). An additional limitation is the potential for circularity if the computational tools evaluated here were used to inform the ClinVar classifications of some of the variants in our calibration set; this impact is likely larger for older tools (e.g., PROVEAN).

Finally, a broader limitation of applying the Tavtigian et al. (2018, 2020) framework is that a lower prior probability of pathogenicity requires higher likelihood ratios to reach evidence levels in both the pathogenic and benign directions. The evidentiary burden is increased symmetrically as a lower prior probability not only requires a higher likelihood ratio to reach the fixed posterior thresholds for pathogenicity, but also requires correspondingly stronger, inverse likelihood ratios to reach the thresholds for benignity. Here, the lower prior for insertions, compared to deletions, made it more difficult for tools to provide additional evidence for benignity, as can be observed by comparing the horizontal lines for insertions and deletions in Figure 3. Ultimately, this resulted in thresholds that may underestimate the strength of the evidence, particularly for benign classification of insertions. However, this conservatism ensures that these thresholds can be applied with high confidence in clinical settings, given the primary objective to avoid overestimating the strength of a piece of evidence.

### Summary

Eight computational tools designed to predict the effects of in-frame insertions and deletions (indels) have been calibrated and systematically evaluated for their applicability in genomic medicine. These predictors demonstrated clear utility in supporting clinical variant classification. The lower strength of evidence they provide relative to established missense variant effect predictors highlights the ongoing need for continued improvement of *in silico* methods for indel variant effect prediction.

## Consortia

The ClinGen Consortium Computational Working Group members include Anne O’Donnell-Luria (co-chair), Predrag Radivojac (co-chair), Moriel Singer-Berk (coordinator), Steven E. Brenner, Alicia B. Byrne, Hannah Carter, Christopher Cassa, Hon-Yun Brian Chung, Sandra T. Cooper, Marc S. Greenblatt, William Hankey, Steven M. Harrison, Kiely James, Vikas Pejaver, Tina Pesaran, Lishuang Shen, Lea M. Starita, and Sean V. Tavtigian.

The ClinGen Consortium Variant Classification Working Group members include Steven M. Harrison (co-chair), Melanie Lacaria (co-chair), Alicia B. Byrne (coordinator), Ahmad Abou Tayoun, Leslie G. Biesecker, Nicole J. Burns, Nadia Carstens, Marina DiStefano, Miranda Durkie, Rajarshi Ghosh, Britt Johnson, Jennifer Johnston, Kristy Lee, Xi Luo, Anne O’Donnell-Luria, Audrey K. O’Neill, Tina Pesaran, Sharon E. Plon, Predrag Radivojac, Heidi L. Rehm, Lea M. Starita, Marcie Steeves, Natasha Strande, and Logan Walker.

## Data and Code Availability

The ClinVar 2023, ClinVar 2025, and gnomAD datasets used are available in Supplementary File 1. Sequence Compressed Reference-orientated Alignment Map (CRAM) files for the RGP dataset are available through dbGaP accession number phs003047 (GREGoR data set). Access is overseen by a data access committee designated by dbGaP and is granted based on the requester’s proposed research use and data use restrictions specified by submitters as defined by consent codes.

## Acknowledgements

This work was supported in part by the NSF GRFP award 2023361442 (H.A.) and the NIH National Human Genome Research Institute (NHGRI) awards U01HG012022 (P.R.), R01HG013350 (V.P.), U24HG007346 (S.E.B.), U01HG011755 (A.O’D.L.). RGP data were provided by Broad Center for Mendelian Genomics with funding by NHGRI grants UM1HG008900 and U01HG011755, by the Chan Zuckerberg Initiative through an advised fund of the Silicon Valley Community Foundation grant 2020-224274, and by research funding from Illumina Inc. ClinGen is primarily funded by the NHGRI with co-funding from the National Cancer Institute (NCI), through the following grants: U24HG009649 (to Baylor/Stanford), U24HG006834 (to Broad/Geisinger), and U24HG009650 (to UNC/Kaiser).

