## Supplementary Information for "Calibration of in-frame indel variant effect predictors for clinical variant classification"

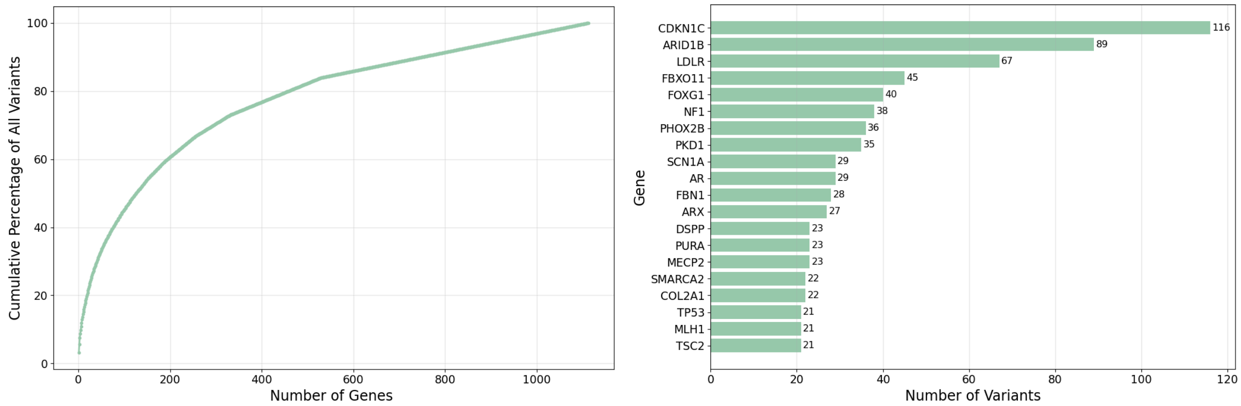


**Supplementary Figure 1.** Distribution of ClinVar 2023 short in-frame indel variants across disease-associated genes. (left) Cumulative percentage of ClinVar 2023 variants accounted for by the top N genes ranked by variant count. (right) Top 20 genes represented in ClinVar 2023 dataset.

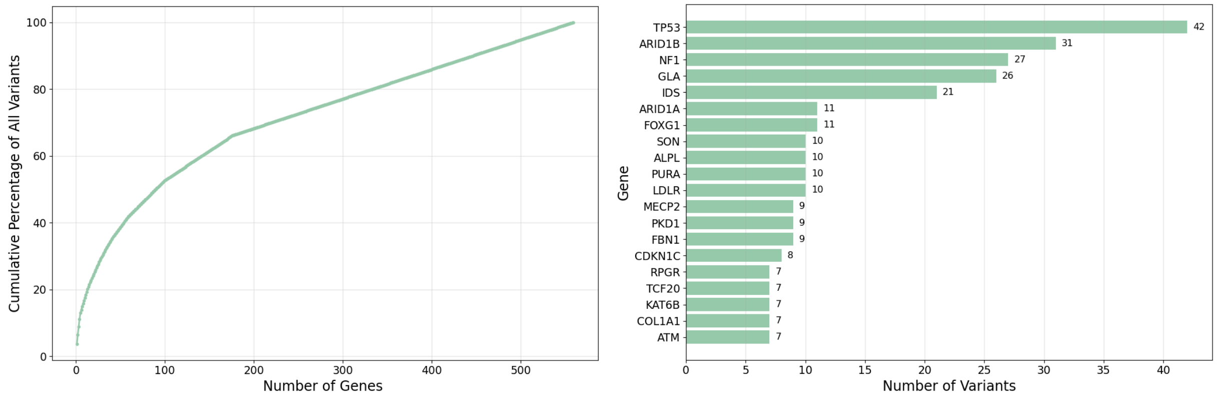


**Supplementary Figure 2.** Distribution of ClinVar 2025 short in-frame indel variants across disease-associated genes. (left) Cumulative percentage of ClinVar 2025 variants accounted for by the top N genes ranked by variant count. (right) Top 20 genes represented in ClinVar 2025 dataset.

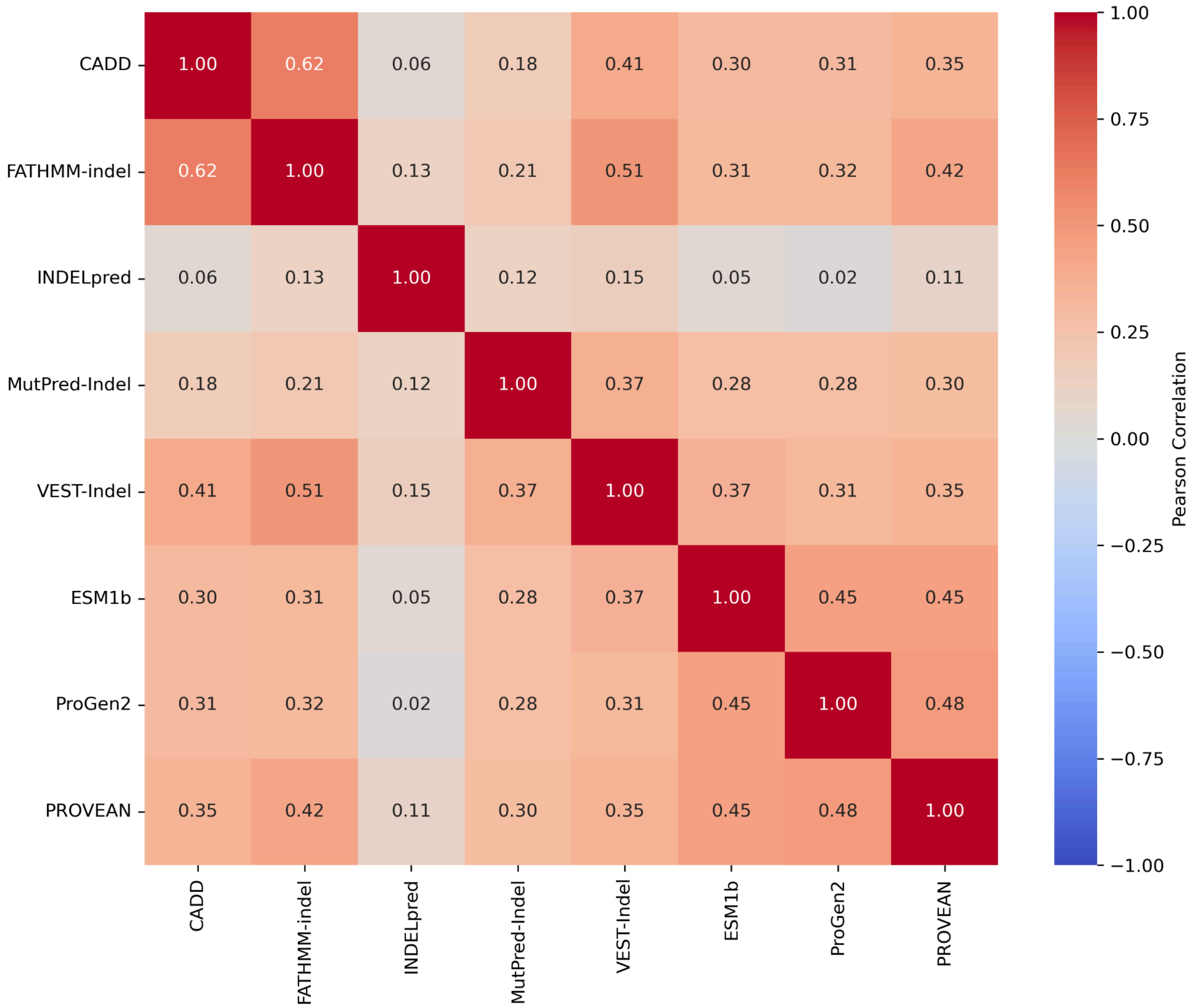


**Supplementary Figure 3.** Pairwise correlation for all methods on gnomAD data set. We calculated the Pearson correlation coefficient for each pair of tools on their intersection of scored variants. Scores were flipped for tools that consider lower scores to be pathogenic (ESM1b, ProGen2, and PROVEAN). We observe notably lower levels of correlation compared to missense variant effect predictors (Pejaver et al., 2022).

| | Variants after removing training data | Variants scored |
|---|---|---|
| CADD | 3625 | 3622 |
| FATHMM-indel | 3593 | 3472 |
| INDELpred | 1783 | 1688 |
| MutPred-Indel | 3017 | 3017 |
| VEST-Indel | 2780 | 2668 |
| ESM1b | 3625 | 3579 |
| ProGen2 | 3625 | 3579 |
| PROVEAN | 3625 | 3576 |

**Supplementary Table 1.** Tools used for analysis. Number of ClinVar 2023 variants provided to each tool for scoring after removing training data and the final number of variants scored for each.

**Deletions**

| | -4 | -3 | -2 | -1 | Indet. | +1 | +2 | +3 | +4 |
|---|---|---|---|---|---|---|---|---|---|
| CADD | | 33 | 197 | 273 | 1860 | 251 | | | |
| FATHMM-indel | 287 | 138 | 194 | 156 | 575 | 388 | 774 | | |
| INDELpred | 148 | 9 | 44 | 31 | 941 | 58 | | | |
| MutPred-Indel | | | 291 | 109 | 1067 | 343 | 146 | 188 | |
| VEST-Indel | | 276 | 247 | 38 | 409 | 206 | 303 | 389 | |
| ESM1b | | | | | 1505 | 487 | 310 | 278 | |
| ProGen2 | | | | 550 | 851 | 323 | 610 | 246 | |
| PROVEAN | 426 | 41 | 240 | 194 | 291 | 979 | 95 | 311 | |

**Insertions**

| | -4 | -3 | -2 | -1 | Indet. | +1 | +2 | +3 | +4 |
|---|---|---|---|---|---|---|---|---|---|
| CADD | | | | 136 | 698 | 144 | 30 | | |
| FATHMM-indel | | | | 112 | 597 | 251 | | | |
| INDELpred | 47 | 0 | 79 | 17 | 314 | | | | |
| MutPred-Indel | | | | 228 | 473 | 121 | 51 | | |
| VEST-Indel | | | | 253 | 337 | 60 | 108 | 42 | |
| ESM1b | | | | | 755 | 79 | 165 | | |
| ProGen2 | | | | | 756 | 116 | 127 | | |
| PROVEAN | | | 71 | 314 | 316 | 36 | 262 | | |

**Supplementary Figure 4.** ClinVar 2023 set variant counts for deletions (top) and insertions (bottom) that met evidence strength level for each tool.

**Deletions**

| | -4 | -3 | -2 | -1 | Indet. | +1 | +2 | +3 | +4 |
|---|---|---|---|---|---|---|---|---|---|
| CADD | | 5 | 40 | 68 | 674 | 105 | | | |
| FATHMM-indel | 41 | 25 | 57 | 37 | 186 | 136 | 375 | | |
| INDELpred | 42 | 3 | 21 | 10 | 714 | 50 | | | |
| MutPred-Indel | | | 66 | 28 | 289 | 155 | 71 | 139 | |
| VEST-Indel | | 56 | 55 | 13 | 198 | 76 | 109 | 194 | |
| ESM1b | | | | | 409 | 206 | 125 | 140 | |
| ProGen2 | | | | 115 | 265 | 110 | 250 | 140 | |
| PROVEAN | 66 | 14 | 62 | 65 | 83 | 372 | 44 | 173 | |

**Insertions**

| | -4 | -3 | -2 | -1 | Indet. | +1 | +2 | +3 | +4 |
|---|---|---|---|---|---|---|---|---|---|
| CADD | | | | 20 | 160 | 40 | 19 | | |
| FATHMM-indel | | | | 9 | 124 | 82 | | | |
| INDELpred | 4 | 1 | 16 | 5 | 187 | | | | |
| MutPred-Indel | | | | 36 | 110 | 42 | 24 | | |
| VEST-Indel | | | | 44 | 71 | 19 | 41 | 14 | |
| ESM1b | | | | | 151 | 28 | 52 | | |
| ProGen2 | | | | | 148 | 26 | 57 | | |
| PROVEAN | | | 9 | 66 | 59 | 11 | 89 | | |

**Supplementary Figure 5.** ClinVar 2025 set variant counts for deletions (top) and insertions (bottom) that met evidence strength level for each tool.

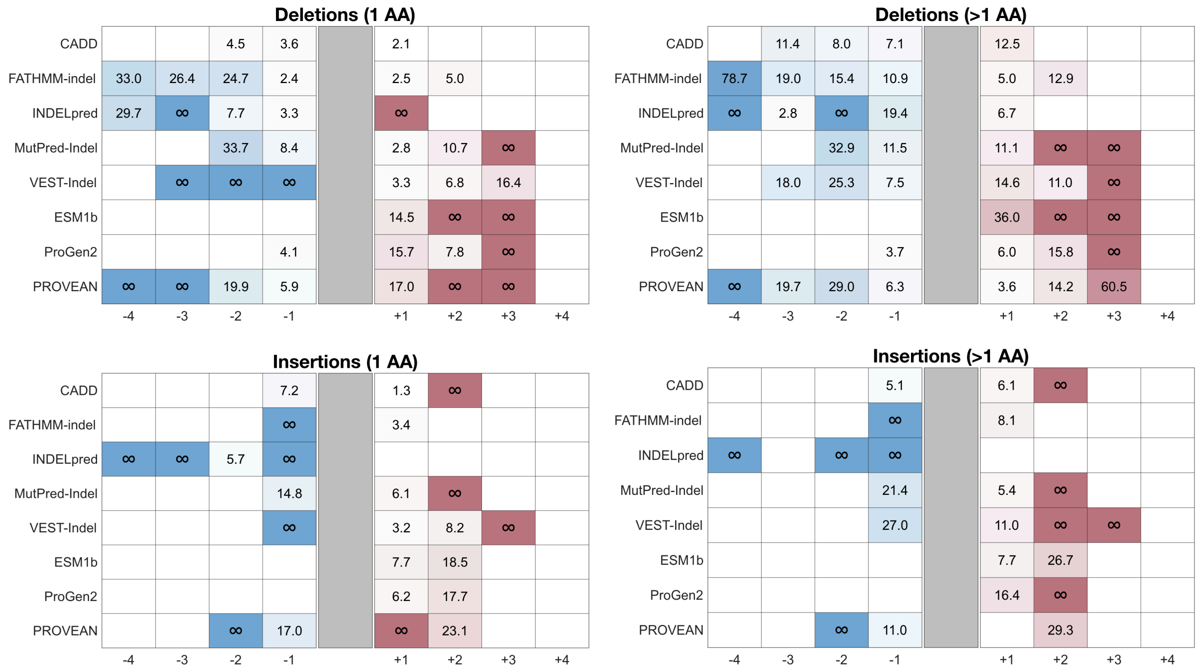


**Supplementary Figure 6.** Likelihood ratios calculated on ClinVar 2025 set calculated separately for shorter (one amino acid change) and longer (greater than one amino acid change and shorter than 17 amino acid change) deletions and insertions. The specific grouping was made to roughly split the available variants in half. The expected likelihood ratios for deletions are 2.39 (supporting, +1), 5.69 (moderate, +2), 13.59 (moderate, +3) and 32.42 (strong, +4). The expected likelihood ratios for insertions are 3.25 (supporting, +1), 10.55 (moderate, +2), 34.28 (moderate, +3) and 111.34 (strong, +4).

| | Deletions | | Insertions | |
|---|---|---|---|---|
| | 1 amino acid | > 1 amino acid | 1 amino acid | > 1 amino acid |
| CADD | 1288 | 1326 | 347 | 661 |
| FATHMM | 1240 | 1272 | 325 | 635 |
| INDELpred | 556 | 675 | 154 | 303 |
| MutPred-Indel | 1012 | 1132 | 298 | 575 |
| VEST-Indel | 834 | 1034 | 256 | 544 |
| ESM1b | 1274 | 1306 | 341 | 658 |
| ProGen2 | 1274 | 1306 | 341 | 658 |
| PROVEAN | 1275 | 1302 | 343 | 656 |

**Supplementary Table 2.** Counts of scored ClinVar 2023 variants per tool for one amino acid indels versus longer indels. The specific grouping into short versus long in-frame indels was made to roughly split the available variants in half.

| | All genes | Disease-associated genes |
|---|---|---|
| | Scored (total 4548 variants) | Scored (total 1140 variants) |
| CADD | 4547 | 1140 |
| FATHMM-indel | 4533 | 1136 |
| INDELpred | 4541 | 1134 |
| MutPred-Indel | 4220 | 1052 |
| VEST-Indel | 4442 | 1133 |
| ESM1b | 4223 | 1053 |
| ProGen2 | 4223 | 1053 |
| PROVEAN | 4314 | 1106 |

**Supplementary Table 3.** Number of rare in-frame indels scored for all genes and disease-associated genes from RGP probands.

**Additional information about indel computational predictors**
Web servers:
CADD: https://cadd.gs.washington.edu/score
VEST-Indel: https://cravat.us/CRAVAT/
PROVEAN: http://provean.jcvi.org/genome_submit_2.php?species=human (accessed via Safari web browser)
FATHMM-indel: http://indels.biocompute.org.uk/
MutPred-Indel: https://mutpred.mutdb.org/mutpredindel/

CADD and VEST-Indel scores are freely available for non-commercial use; commercial uses require a license. FATHMM-indel is freely available via the web server at the link above. MutPred-Indel allows free use of the web site for non-commercial and commercial use, with the number of variants per query limited to 100. MutPred-Indel source code and software are freely available for non-commercial use and require a license for commercial use. The PROVEAN web server was accessed at the link above without registration using the Safari web browser; the server was operational at the time of analysis despite being listed as retired by JCVI. The standalone PROVEAN software remains available for download under the GNU General Public License v3. INDELpred, ESM1b, ProGen2, and MutPred-Indel were run locally using code downloaded from their respective GitHub repositories or website for MutPred-Indel. ESM1b is released under the MIT License and ProGen2 under the BSD-3-Clause License. INDELpred does not specify an explicit code license.
